# A smooth muscle-like niche facilitates lung epithelial regeneration

**DOI:** 10.1101/565085

**Authors:** Alena Moiseenko, Ana Ivonne Vazquez-Armendariz, Xuran Chu, Stefan Günther, Kevin Lebrigand, Vahid Kheirollahi, Susanne Herold, Thomas Braun, Bernard Mari, Stijn De Langhe, Chengshui Chen, Xiaokun Li, Werner Seeger, Jin-San Zhang, Saverio Bellusci, Elie El Agha

## Abstract

The mammalian lung is a highly complex organ due to its branched, tree-like structure and diverse cellular composition. Recent efforts using state-of-the-art genetic lineage tracing and single-cell transcriptomics have helped reduce this complexity and delineate the ancestry and fate of various cell subpopulations during organogenesis, homeostasis and repair after injury. However, mesenchymal cell heterogeneity and function in development and disease remain a longstanding issue in the lung field. In this study, we break down smooth muscle heterogeneity into the constituent subpopulations by combining *in vivo* lineage tracing, single-cell RNA sequencing and *in vitro* organoid cultures. We identify a repair-supportive mesenchymal cell (RSMC) population that is distinct from pre-existing airway smooth muscle cells (ASMC) and is critical for regenerating the conducting airway epithelium. Progenitors of RSMCs are intertwined with airway smooth muscle, undergo active WNT signaling, transiently acquire the expression of the smooth muscle marker ACTA2 in response to epithelial injury and are marked by PDGFRα expression. Our data simplify the cellular complexity of the peribronchiolar domain of the adult lung and represent a forward step towards unraveling the role of mesenchymal cell subpopulations in instructing epithelial behavior during repair processes.

## Introduction

The respiratory tract consists of a conducting zone as well as a transitional and respiratory zone. While the latter zone represents the site of gas exchange between inhaled air and the pulmonary circulation, the conducting zone represents the first line of defense and domain of contact between the respiratory system and the external environment. As such, the trachea, bronchi, bronchioles and terminal bronchioles, constituting the conducting zone, are targets for a myriad of airborne pathogens, pollutants, toxins and other irritants.

Club cells, previously known as exocrine bronchiolar cells or Clara cells (Clara, 1937; Winkelmann and Noack, 2010), are dome-shaped, non-ciliated secretory cells that populate the bronchiolar region of the lung epithelium. These cells can be identified by the expression of secretoglobin family 1A member 1 (SCGB1A1, a.k.a. Clara cell 10 KDa secretory protein CC10 or CCSP). Besides playing a major role in detoxifying the airways through biotransformation of inhaled xenobiotics, club cells also serve as long-term progenitors for ciliated and secretory cells in the bronchioles (Rawlins et al., 2009; Stripp et al., 1995). Owing to significant expression levels of cytochrome P450 family 2 subfamily f polypeptide 2 (CYP2F2), club cells are selectively targeted by naphthalene; they metabolize it into a toxic derivative and consequently undergo necrosis and depletion (Boyd, 1977; Fanucchi et al., 1997; Mahvi et al., 1977). Interestingly, surviving variant club cells (CYP2F2^-^) located within neuroepithelial bodies (NEBs) and at bronchioalveolar duct junctions (BADJs) mediate the repair process (Giangreco et al., 2002; Hong et al., 2001; Reynolds et al., 2000). Accordingly, the naphthalene injury model is widely used in experimental mice to study mechanisms of club cell replenishment and, in a broader sense, epithelial regeneration.

During embryonic lung development, epithelium-derived WNT ligand, WNT7b, acts on adjacent airway and vascular smooth muscle cell (ASMC and VSMC, respectively) progenitors to induce platelet-derived growth factor receptor alpha and beta (*Pdgfra* and *Pdgfrb*) expression (Cohen et al., 2009). The effect of β-catenin (CTNNB1)-mediated WNT signaling on SMC differentiation has been shown to be mediated by the extracellular matrix protein tenascin C (TNC) (Cohen et al., 2009). Therefore, the WNT/TNC/PDGFR pathway is a key signaling event for SMC formation in the developing lung. Moreover, early ASMC precursors can be identified by the expression of fibroblast growth factor 10 (*Fgf10*) (El Agha et al., 2014; Mailleux et al., 2005), a key developmental gene for lung branching morphogenesis and maintenance of epithelial progenitor cells during embryonic lung development (Bellusci et al., 1997a; Ramasamy et al., 2007; Sekine et al., 1999).

Previous studies have implicated similar epithelial-mesenchymal interactions in progenitor cell activation after injury in the adult lung. In particular, it has been shown that following naphthalene injury in adult mice, the airway epithelium-derived WNT ligand, WNT7b, leads to upregulation of *Fgf10* in ASMCs. FGF10, in turn, acts on surviving club cells expressing FGF receptor 2-IIIb (*Fgfr2b*) to induce epithelial regeneration (Volckaert et al., 2011). In another elegant study, single-cell RNA sequencing (scRNA-seq), organoid assays and genetic cell ablation *in vivo* and *in vitro* have shown that leucine-rich repeat-containing G-protein coupled receptor 6 (*Lgr6*) expression identifies an ASMC subpopulation that promotes epithelial repair after naphthalene injury in a WNT-FGF10-mediated fashion (Lee et al., 2017). Therefore, there is a notion that the peribronchiolar region represents a mesenchymal niche for epithelial stem/progenitor cells and facilitates the repair process after injury.

In this study, we took advantage of the temporal control of CreERT2 activity to lineage-trace mesenchymal cells at different stages of homeostasis, injury and repair. We performed scRNA-seq, to investigate the heterogeneity of labeled mesenchymal cells, and organoid assays to test their ability to support club-cell growth *in vitro*. Remarkably, we found that pre-existing ASMCs are not the sole niche cells for surviving club cells after naphthalene injury as previously described. We identify a novel mesenchymal population, termed repair-supportive mesenchymal cells (RSMCs), that transiently expresses the smooth muscle cell marker, alpha smooth muscle actin (*Acta2*), invades the ASMC sheet and contributes to the repair process. Our data highlight mesenchymal cell heterogeneity even within a confined region like the peribronchiolar space and demonstrate dynamism of mesenchymal cells in response to epithelial injury.

## Materials and Methods

### Mice and tamoxifen administration

*Acta2-CreERT2 (STOCK_Tg(Acta2-cre/ERT2)12Pcn)* mice (Wendling et al., 2009) were kindly provided by Dr. Pierre Chambon (University of Strasbourg, France). *Fgf10^flox^ (B6;129-Fgf10tm1.2Sms/J)* mice (Abler et al., 2009; Urness et al., 2010) were a generous gift from Dr. Suzanne L. Mansour (University of Utah, USA). The mouse strains *tdTomato^flox^* (*B6;129S6-Gt(ROSA)26Sor^tm9(CAG-tdTomato)Hze^/J*, stock number 007905), *Ctnnb1^flox^* (*B6.129-Ctnnb1tm2Kem/J*, stock number 004152)*, Gli1^CreERT2^* (*STOCK_Gli1^tm3(cre/ERT2)Alj^*, stock number 007913) and *Pdgfra^H2B-EGFP^* (B6.129S4-*Pdgfra^tm11(EGFP)Sor^*/J) were purchased from the Jackson laboratory. *Scgb1a1^CreERT2^ (B6N.129S6(Cg)-Scgb1a1tm1(cre/ERT)Blh/J)* mice crossed with *tdTomato^flox^* (*B6.Cg-Gt(ROSA)26Sortm14(CAG-tdTomato)Hze/J)* were kindly provided by Prof. Christos Samakovlis (Justus-Liebig University Giessen). To induce CreERT2-mediated genetic recombination, *Acta2-CreERT2* and *Gli1^CreERT2^* mice were fed tamoxifen-containing food (0.4 g of tamoxifen per kg of food) (Altromin, Germany). Tamoxifen powder was purchased from Sigma-Aldrich. Animal experiments were approved by the Regierungspraesidium Giessen (2/2016) and University of Alabama at Birmingham Institutional Animal Care and Use Committee (IACUC).

### Naphthalene treatment

For naphthalene treatment, 8-12-week-old mice were used. Naphthalene (Sigma) was dissolved in corn oil (20 mg/mL) and was administered intraperitoneally at a dose of 0,275 mg naphthalene per g of body weight.

### Immunofluorescence

For histological analysis, lungs were perfused with PBS and fixed with 4% paraformaldehyde. Then, tissues were embedded in paraffin and cut into 5 μm-thick sections. Immunofluorescence (IF) staining was performed using monoclonal anti-ACTA2 (Sigma, 1:200), monoclonal anti-CC10 (Santa Cruz Biotechnology, 1:200), monoclonal anti-CD45 (Novus Biologicals, 1:200) and monoclonal anti-E-cadherin (BD Biosciences, 1:200) antibodies. Staining for CD45 and E-cadherin was performed after antigen retrieval with citric buffer and boiling for 15 min. The latter stainings were followed by staining with polyclonal anti-tdTomato (anti-RFP) antibodies (Rockland, 1:200) in order to recover the tdTomato signal. Nuclei were counterstained with DAPI (4’,6-diamidino-2-phenylindole) (Life Technologies). Fluorescent images were acquired using Leica DM550 B fluorescence microscope (equipped with Leica DFC360 FX camera) or an upgraded version of a TCS SP5 confocal microscope (both from Leica Microsystems). For quantitative analysis of immunofluorescence, multiple images were used (n > 8). For each experiment, sections from at least 3 independent lungs were analyzed. Tile scans were obtained using EVOS FL Auto 2 Cell Imaging system (Invitrogen). For imaging of thick *Gli1^CreERT2^, tdTomato^flox^* samples, lungs were perfused with 1% low-melting agarose and cut into 200 μm-thick sections using a vibratome (Leica Microsystems). Sections were then fixed and stained with ACTA2 similarly to paraffin-embedded sections and imaged by confocal microscopy. Three-dimensional (3D) reconstruction of z-stacks was performed using LAS X software (Leica Microsystems).

### Flow cytometry analysis and cell sorting

Lungs were isolated in Hank’s balanced salt solution (HBSS, Gibco), cut into small pieces with a sharp blade and treated with 0.5% collagenase type IV in HBSS (Life technologies) at 37°C for 45 minutes. Lung homogenates were passed through 18, 21 and 24G needles and then through 70 and 40 μm cell strainers (BD Biosciences) to obtain single-cell suspensions. Cells were centrifuged at 4°C at 1000 rpm for 5 min and then resuspended in staining solution (0,1% sodium azide, 5% FCS in PBS) containing anti-PDGFRα antibodies (APC-conjugated, Biolegend, 1:100), anti-EpCAM (APC-Cy7-conjugated, Biolegend, 1:100), anti-CD31 (Pacific Blue-conjugated, Biolegend, 1:100) and anti-CD45 (Pacific Blue-conjugated, Biolegend, 1:100) for 20 min on ice in the dark. Then, cells were washed with 0.1% sodium azide in PBS. Flow cytometry measurements of labeled cells and cell sorting were done using FACSAria III cell sorter (BD Biosciences).

### Quantitative Real-time PCR

For bulk RNA extraction, accessory lobes were lyzed and homogenized using Bullet Blender Blue (Next Advance, USA), and RNA was extracted using RNeasy kit (Qiagen, Germany). Extraction of RNA from sorted cells was performed using RNeasy Micro kit (Qiagen). Primers were designed using the Universal Probe Library Assay Design center program (Roche Applied Science). Quantitative real-time PCR was performed using Light Cycler 480 II (Roche Applied Science). Hypoxantine-guanine phosphoribosyltransferase (*Hprt*) was used to normalize gene expression. Data were presented as mean expression relative to *Hprt* ± sem.

### Single-cell RNA sequencing

Single-cell RNA-seq was performed using the ICELL8 platform (Takara Bio) using the V2 protocol with in-chip preamplifcation. Samples were pre-sorted into HBSS (Gibco) at a concentration of 20 cells/µL and stained with Hoechst dye (Invitrogen) for 5 min at RT prior to loading onto the MSND dispenser. For each sample 3 dispenses were made on one 5184 well chip resulting in 348 and 459 wells containing single cells after selection in CellSelect software (Takara Bio). Selected wells were processed for RT, in-chip preamplification and cDNA was pooled for final library preparation using standard Nextera protocol (Illumina). Library was checked in Labchip Gx touch 24 (Perkin Elmer) and sequenced on Nextseq500 (Illumina) using V2 chemistry and paired-end setup (11bp Read1 for cell barcode and 80bp Read2 for cDNA template sequencing). A total of 537M reads were obtained.

### Single-cell data analysis

Complementary DNA (cDNA) reads were trimmed for poly(A) tails using BBMAP (unpublished, https://jgi.doe.gov/data-and-tools/bbtools/). Reads of length under 50 bases were discarded. The 456M remaining reads were then aligned using STAR (release v2.4.0a) against *Mus musculus* build mm10 following ENCODE RNA-seq mapping recommendations. STAR indices were generated using Ensembl GTF file (release 83). For counting based on read counts, we used the Dropseq Core Computational Protocol version 1.12 (dropseq.jar) (Macosko et al., 2015). Gene to cell count table matrix was then used for statistical analysis performed within the Seurat package (http://satijalab.org/seurat/) (Butler et al., 2018). The pipeline goes through cell and gene filtering, data normalization, then finding the most variable genes to perform principal component analysis (PCA) and t-distributed stochastic neighbor embedding (t-SNE) dimension reduction for visualization purpose. We first created a Seurat object including all cells and all genes present in at least 5 different cells. We then filtered out cells with more than 95% dropouts or less than 50.000 read counts, we also discard cells with more than 20% of counts from mitochondrial genes or 10% counts from ribosomal genes. We then normalized the count values for the 305 remaining cells to a scaling factor corresponding to the median read counts per cell. We next identified most variable genes by plotting genes into bins based on X-axis (average expression) and Y-axis (dispersion) using cutoffs X = 3 and Y = 0.1, identifying 1,212 genes. We then performed a PCA, after scaling and centering of the data across these genes. This ensured robust identification of the primary structures in the data. We identified the first 8 principal components as statistically significant and use it for t-SNE calculation (perplexity=30). The t-SNE procedure returns a two dimensional embedding of single cells utilized for dataset visualization. Cells with similar expression signatures of genes within the variable set, and therefore similar PC loadings, are most likely to localize near each other in the embedding, and hence distinct cell types form two-dimensional point clouds across the t-SNE map. Identification of the clusters of cells was determined using the FindClusters Seurat function (resolution=1). The Seurat FindAllMarkers function was used to identify differentially expressed genes across each cluster (min.pct = 0.25, thresh.use = 0.25)

### Organoid assay

Sorted mesenchymal cells (tdTom+ PDGFRα+, tdTom+ PDGFRα^-^ and tdTom+) from *Acta2-CreERT2; tdTomato^flox^* mice and epithelial cells (tdTom+) from *Scgb1a1^CreERT2^; tdTomato^flox^* mice were centrifuged and resuspended in cell culture medium (Dulbecco’s Modified Eagle Medium, Life Technologies). Concentrations were adjusted to 1×10^3^ epithelial cells in 25 μL media and 2×10^4^ mesenchymal cells in 25 μL media per insert (12 mm cell culture inserts with 0.4 µm membrane (Millipore) were used). Epithelial and mesenchymal cell suspensions were mixed and cold Matrigel^®^ growth factor-reduced (Corning) at a 1:1 dilution was added resulting in 100 μL final volume per insert. Matrigel cell suspensions were added on top of the filter membrane of the insert and incubated at 37°C for 5 min. Then 350 μL of medium was added per well. Cells were incubated under air-liquid conditions at 37°C with 5% CO_2_ for 2 weeks. Media were changed 3 times per week.

### Statistical analysis and figure assembly

Quantitative data were assembled using GraphPad Prism software (GraphPad Software) and presented as average values +/-sem. Statistical analyses were performed using one-way ANOVA unless stated otherwise, and the number of biological replicates is indicated in the corresponding figure legends. Results were considered significant if P<0.05. Figures were assembled using Adobe Photoshop and Illustrator (CS5/6).

## Results

### Genetic deletion of *Ctnnb1* or *Fgf10* in pre-existing ASMCs does not compromise epithelial regeneration

The cell turnover rate in the lung is generally low under homeostatic conditions, particularly in terminally differentiated cells such as ASMCs. It has been proposed that activation of canonical CTNNB1-mediated WNT signaling in ASMCs is critical for inducing epithelial repair following naphthalene injury. In order to investigate whether pre-existing, terminally differentiated ASMCs mediate the repair process, *Acta2-CreERT2; Ctnnb1^flox/flox^* (experimental) and *Ctnnb1^flox/flox^* (control) mice were fed tamoxifen-containing pellets for 2 weeks, followed by a washout period of 2 weeks, and then treated with an intraperitoneal (IP) injection of naphthalene or corn oil (Fig. S1A). Animals were sacrificed at day 14 after injury. Body weight was monitored on a daily basis and the rate of weight loss and gain was virtually indistinguishable between the experimental and control groups, showing a typical 20% weight loss at day 3 followed by gradual recovery up to day 14 (Fig. S1B). Immunofluorescence for SCGB1A1 showed similar levels of immunoreactivity in control and experimental animals at day 14 (Fig. S1C, D). Moreover, qPCR on lung homogenates showed comparable expression levels for the repair markers *Scgb1a1*, *Wnt7b* and *Fgf10* in both groups (Fig. S1E).

A similar experimental setup was applied to *Acta2-CreERT2; Fgf10^flox/flox^* mice (Fig. S1F) and the effect of genetic deletion of *Fgf10* in pre-existing ASMCs was negligible in terms of weight loss/gain (Fig. S1G), SCGB1A1 immunoreactivity (Fig. S1H, I) and expression levels of repair markers (Fig. S1J). As such, these data indicate that canonical WNT signaling and FGF10 production in pre-existing ASMCs are dispensable for club cell replenishment following naphthalene injury.

### WNT signaling and FGF10 production in ACTA2+ cells targeted after injury are critical for epithelial regeneration

We next set out to determine whether CTNNB1-mediated WNT signaling is required for recovery after naphthalene injury. *Acta2-CreERT2; Ctnnb1^flox/flox^* (experimental) and *Ctnnb1^flox/flox^* (control) mice received an IP injection of naphthalene or corn oil and were then fed tamoxifen-containing pellets throughout the whole duration of injury and recovery (Fig. 1A). This experimental setup allowed genetic deletion of *Ctnnb1* not only in *bona fide* ASMCs but also in potential newly recruited cells that transiently acquired *Acta2* expression following naphthalene injury. Lungs were harvested at multiple time points up to day 14 (Fig. 1A). Intriguingly, experimental mice displayed more severe weight loss in response to injury as well as suboptimal recovery compared to controls (Fig. 1B). Quantification of SCGB1A1 immunofluorescence showed that although the expression patterns were comparable at the peak of injury (day 3), experimental mice revealed progressive impairment in club cell replenishment up to day 14 (Fig. 1C, D). Quantitative real-time PCR for *Scgb1a1* using lung homogenates confirmed the histological observations and also revealed decreased *Wnt7b* and *Fgf10* expression levels in experimental animals compared to controls (Fig. 1E). Immunofluorescence for KI67, which marks all phases of the cell cycle except for G_0_ phase, showed a 2-fold decrease in immunoreactivity in the airway epithelium upon loss of β-catenin, indicating less cell-cycle progression (Fig. S2). A similar analysis was carried out on *Acta2-CreERT2; Fgf10^flox/flox^* mice and the results showed similar detrimental consequences for *Fgf10* loss of function during the repair process (Fig. 1F-J). Therefore, we conclude that WNT signaling and FGF10 production in ACTA2+ cells, arising after naphthalene injury, are important for the repair process.

**Figure 1:**
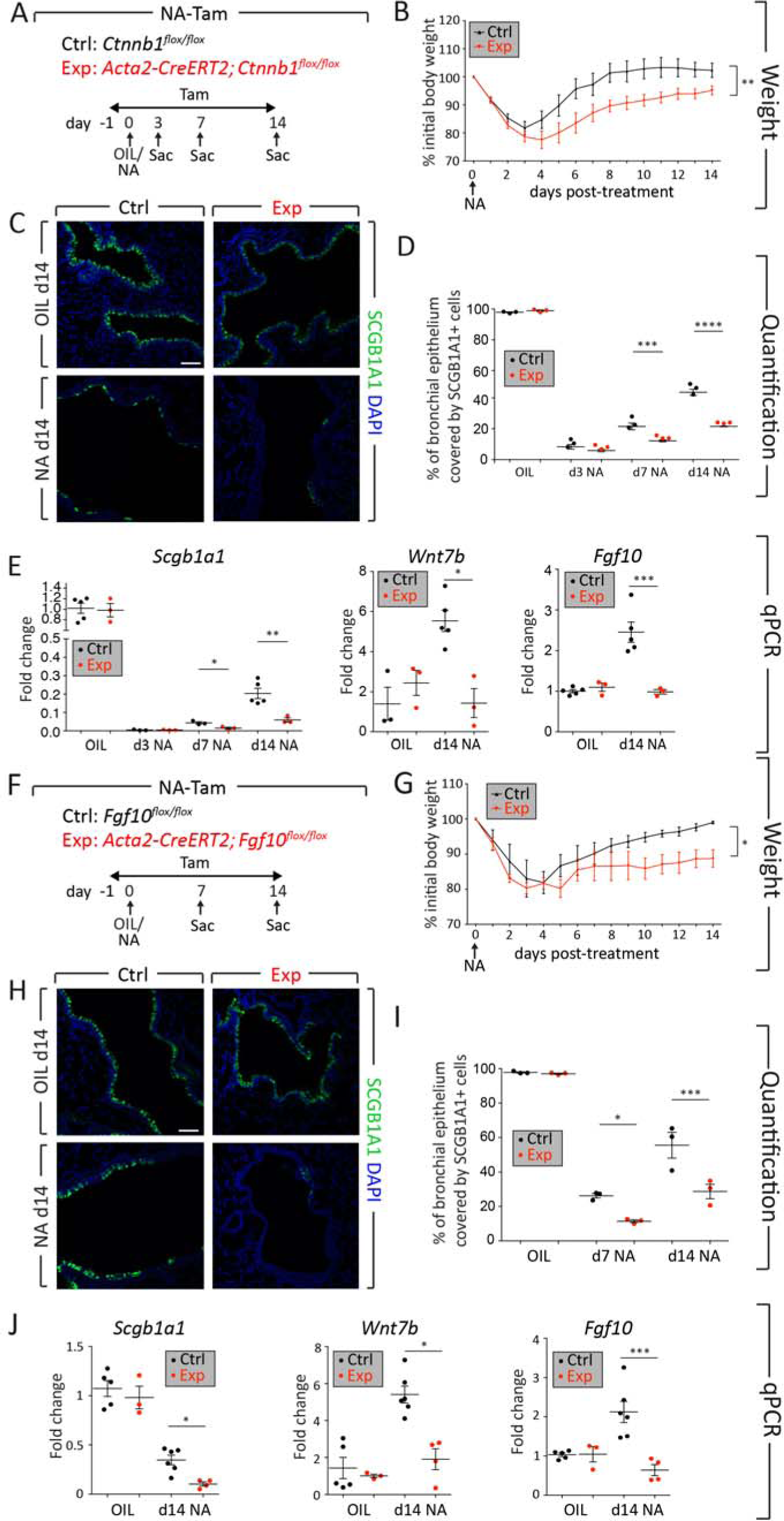
Deletion of *Ctnnb1* or *Fgf10* in ACTA2+ cells after naphthalene injury impairs epithelial regeneration. (A, F) Experimental setup and time line of tamoxifen and naphthalene treatment. (B, G) Weight loss curve showing more severe weight loss in experimental mice. (C, H) Immunofluorescence for SCGB1A1 on control and experimental lungs and (D, I) quantification of immunofluorescence, demonstrating decreased abundance of SCGB1A1+ cells in the lungs of experimental mice compared to controls. (E, J) qPCR on lung homogenates showing decreased levels of *Scgb1a1*, *Wnt7b* and *Fgf10* expression in experimental mice compared to controls. Ctrl – Control mice, Exp – Experimental mice, NA – Naphthalene, Sac – Sacrifice, Tam – Tamoxifen. Scale bars: 50 μm in C; 100 μm in H. N=3 for all experiments except for N=5 for Ctrl-OIL and N=5 for Ctrl-NA d14 in B and E; N = 3 for each experiment in C, D, H and I; N=5 for Ctrl-OIL, N=3 for Exp-OIL, N=6 for Ctrl-NA d14 and N=4 for Exp-NA d14 in G and J. * P<0.05, ** P<0.01, *** P<0.001, **** P<0.0001.

### Mesenchymal cells transiently expressing *Acta2* are recruited during epithelial injury and repair

In order to investigate whether an unknown mesenchymal population transiently acquires *Acta2* expression in response to club cell depletion, the *Acta2-CreERT2; tdTomato^flox^* lineage-tracing tool was employed. Animals were randomly allocated into three groups receiving either tamoxifen food prior to corn oil injection (Tam-Oil), tamoxifen food prior to naphthalene injection (Tam-NA) or a naphthalene injection followed by tamoxifen food (NA-Tam) (Fig. 2A). Animals were sacrificed at day 14 after naphthalene or corn oil injections. Such approach allowed labeling steady-state ASMCs (Tam-Oil), pre-existing ASMCs during repair after injury (Tam-NA) and total ACTA2+ cells during repair after injury (pre-existing ASMCs and newly formed SMCs or newly recruited SMC-like ACTA2+ cells) (NA-Tam). As expected, the lineage label, tdTomato, colocalized with the ACTA2 stain in the Tam-Oil (Fig. 2B-E) and Tam-NA groups (Fig. 2F-I). Intriguingly, imaging of NA-Tam lungs revealed, in addition to ACTA2+ tdTomato+ cells in the ASMC layer, the presence of ACTA2− tdTomato+ cells that were intertwined with ASMCs (Fig. 2J-M). In some instances, these ACTA2− tdTomato+ cells were observed between the ASMC layer and the airway epithelium (Fig. 2M). The presence of ACTA2^-^ tdTomato+ cells indicated that those cells were descendants of a mesenchymal population that transiently acquired *Acta2* expression after naphthalene injury. The number of lineage-labeled cells at day 14 was quantified by flow cytometry and the results showed that while there was no significant change between the Tam-Oil and Tam-NA groups, the NA-Tam group revealed a significant increase in tdTomato+ cell count compared to the other two groups (Fig. 2N). Further analysis of the dynamics of tdTomato+ cells in NA-Tam lungs showed a significant increase in the abundance of tdTomato+ cells from day 3 to days 7/14 after naphthalene injury (Fig. 2P). Interestingly, some tdTomato+ cells located in the peribronchiolar space gradually lost *Acta2* expression over time (Fig. 2Q-S). Therefore, we conclude that after naphthalene injury, a mesenchymal population that is distinct from pre-existing ASMCs transiently acquires *Acta2* expression and is potentially involved in the repair process.

**Figure 2:**
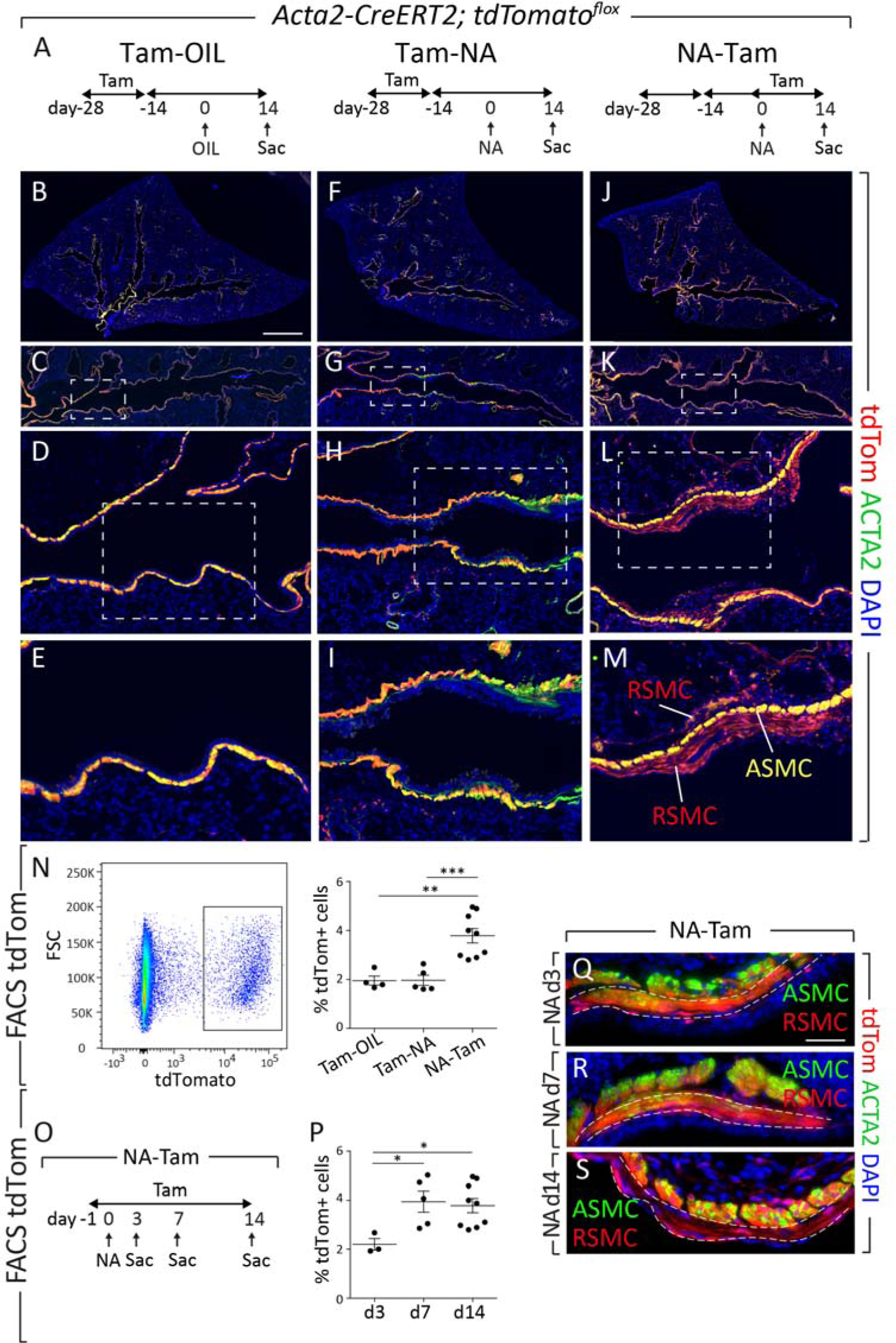
A population of mesenchymal cells transiently expressing ACTA2 is recruited to the sites of airway repair. (A) Experimental setup and time line of tamoxifen and naphthalene/corn oil treatment. (B, F, J) Tile scans of left lobes, (C, G, K) higher magnification of the conducting airway, (D, H, L) higher magnification of the dashed boxes in C, G, K and (E, I, M) high magnification of the dashed boxes in D, H, L are shown for control oil-treated mice (Tam-OIL), naphthalene-treated mice where pre-existing SMCs were labeled (Tam-NA) and naphthalene-treated mice where pre-existing and newly formed ACTA2+ cells were labeled (NA-Tam). Note the appearance of an additional population of ACTA2− tdTomato+ cells (RSMC) in the NA-Tam group (M). (N) Flow cytometry analysis showing an increase in the percentage of tdTomato+ cells in NA-Tam group compared to Tam-OIL and Tam-NA groups. (O) Experimental setup and time line of naphthalene and tamoxifen treatment for analysis of different time points for the NA-Tam group. (P) Flow cytometry analysis demonstrating an increase in the percentage of tdTomato+ cells from d3 to d7 and d14. (Q-S) Immunofluorescence for ACTA2 on NA-Tam lungs collected at d3, d7 and d14. tdTomato and ACTA2 colocalize in RSMCs at d3 and d7 but not at d14. ASMC – Airway smooth muscle cells, FSC – Forward scatter, RSMC – Repair-supportive mesenchymal cells. Scale bars: 1400 μm in B, F, J; 730 μm in C, G, K; 100 μm in D, H, L; 25 μm in E, I, M; 33 μm in Q, R, S. N=4 for Tam-OIL, N=5 for Tam-NA, N=9 for NA-Tam in B-N; N=3 for d3 NA-Tam, N=5 for d7 NA-Tam, N=9 for d14 NA-Tam in P-S. * P<0.05, ** P<0.01, *** P<0.001.

### Single-cell analysis reveals a mesenchymal niche that is associated with epithelial regeneration

We next performed scRNA-seq on sorted, lineage-traced tdTomato+ cells in order to delineate the complexity and heterogeneity of these cells and to identify emerging cell populations that are potentially involved in the repair process in NA-Tam samples. Bulk tdTomato+ cells from Tam-NA (n=3 animals) and NA-Tam (n=3 animals) were used to dispense single cells onto an ICELL8 chip (Fig. 3). Following library preparation and RNA-seq, quality control yielded 122 cells from the Tam-NA group and 183 cells from the NA-Tam group (Figs. 3A-C). The median reads per cell was 303,934 and the median number of genes detected per cell was 1,435 (Fig. S3A, B). Bioinformatic analysis revealed the presence of 5 cellular clusters corresponding to ASMCs/VSMCs, endothelial cells, macrophages and 2 additional clusters that showed typical matrix fibroblast signatures (Fig. 3A-C), according to Lung Gene Expression iN Single-cell (LungGENS) database (https://research.cchmc.org/pbge/lunggens/). A recent report using a Drop-seq scRNA-seq platform (10x Genomics) to analyze whole lung mesenchyme from uninjured and bleomycin-challenged lungs revealed the presence of two matrix fibroblast subpopulations that could be discriminated by the expression of either *Col13a1* and *Col14a1* (Xie et al., 2018). Our two additional clusters displayed partial overlap with these two matrix fibroblast subpopulations. Importantly, analysis of Tam-NA samples showed distribution of tdTomato+ cells between ASMCs/VSMCs (41%), COL14A1+ matrix fibroblast-like cells (15%), endothelial cells (15%), macrophages (4%) and COL13A1+ matrix fibroblast-like cells (25%) (Fig. 3B, E). On the other hand, analysis of NA-Tam samples revealed distribution of tdTomato+ cells between ASMCs/VSMCs (14%), COL14A1+ matrix fibroblast-like cells (19%), endothelial cells (3%), macrophages (5%) and COL13A1+ matrix fibroblast-like cells (59%) (Fig. 3C, F). Therefore, there was a significant emergence of the COL13A1+ matrix fibroblast-like population in NA-Tam lungs. Since deletion of *Ctnnb1* and *Fgf10* only in NA-Tam but not in Tam-NA led to repair impairment (Fig. 1 and Fig. S1), we decided to call this emerging population a repair-supportive mesenchymal cell (RSMC) population. Among the top enriched genes in this cell population was *Pdgfra* (Fig. 3G, S3C). In order to validate these data, *Pdgfra^H2B-EGFP^* were also subjected to naphthalene injury and lungs were examined at day 7 after injury (Fig. S3I). Nuclear EGFP was indeed detected in ACTA2-cells in the peribronchiolar region (Fig. S3I). The total number of differentially expressed genes in the RSMC cluster was 99 genes including *Tcf21, Npnt, Pdgfra, Lbh* and *Igfbp6* (Fig. 3G, S3J). Out of the 99 genes that were enriched in RSMCs, 54 genes were found to be expressed in the previously reported COL13A1+ matrix fibroblasts (Xie et al., 2018) while the rest 45 genes were uniquely expressed in the RSMC cluster.

**Figure 3:**
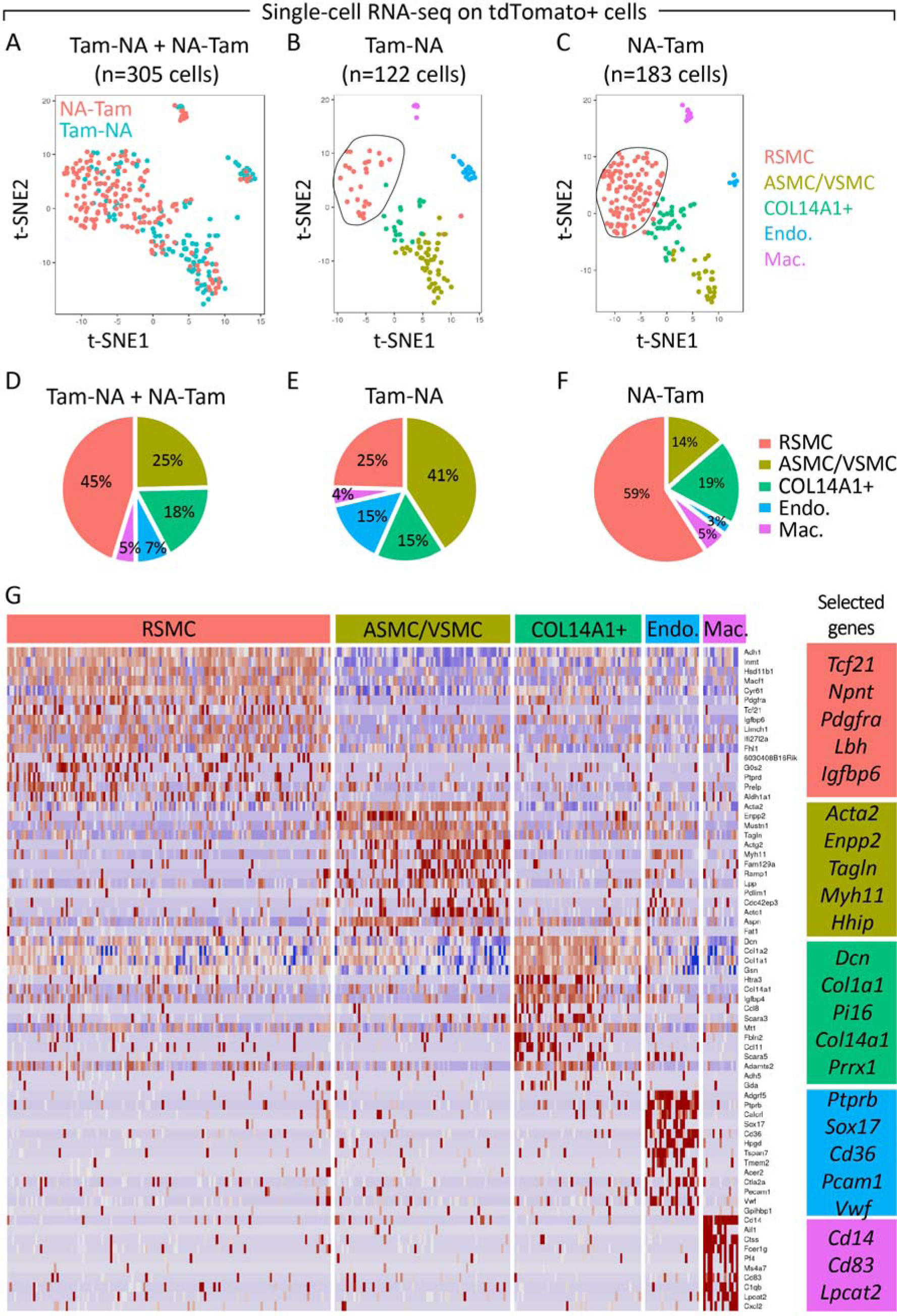
Single-cell RNA sequencing of tdTomato+ cells isolated from Tam-NA and NA-Tam groups identifies a novel cell population involved in epithelial repair. (A) t-SNE plot showing five identified cellular clusters – ASMC/VSMC, COL14A1+ matrix fibroblasts, endothelial cells, macrophages and RSMC. (B) t-SNE plot showing the distribution of tdTomato+ cells from Tam-NA lungs among the identified clusters. Note the low relative number of cells in the RSMC cluster. (C) t-SNE plot showing the distribution of tdTomato+ cells from NA-Tam lungs among the identified clusters. Note the higher abundance of cells in the RSMC cluster. (D-F) Pie charts showing the relative distribution of tdTomato+ cells among the various cellular clusters. (G) Heat map showing the top highly enriched genes in the different clusters. Groups of selected genes are shown in the corresponding boxes. N=3 (pooled) for both Tam-NA and NA-Tam experiments.

Additionally, cells within the ASMC/VSMC subpopulation revealed a total of 139 differentially expressed genes and displayed high expression levels for typical SMC markers such as *Acta2, Enpp2, Tagln, Myh11* and *Hhip* (Fig. 3G, S3E). COL14A1+ matrix fibroblast-like cells revealed 67 differentially expressed genes including *Dcn, Pi16, Col14a1* and *Prrx1* (Fig. 3G, S3F). Out of the 67 genes that were enriched in our COL14A1+ matrix fibroblasts, 40 genes were expressed in the previously reported COL14A1+ matrix fibroblasts (Xie et al., 2018). Cells within the endothelial cluster displayed a total of 223 differentially expressed genes and enrichment in the expression of typical endothelial markers such as *Sox17, Pcam1* and *Vwf* (Fig. 3G, S3G). Last but not least, macrophages showed a total of 201 differentially expressed genes with high expression levels of *Cd14, Cd83, Lpcat2, Il1b* and others (Fig. 3G, S3H).

### PDGFRα expression discriminates between RSMCs and SMCs during lung repair

We then decided to analyze the dynamics of the tdTomato+ PDGFRα+ cell population that emerged in the NA-Tam group. Flow cytometry was carried out at days 3, 7 and 14 following naphthalene injection (Fig. 4A, B). The percentage of tdTomato+ PDGFRα+ out of total tdTomato+ cells increased steadily from day 3, to day 7 and then day 14 (Fig. 4C). qPCR on sorted cells revealed higher expression levels for *Pdgfra*, *Fgf10*, glioma-associated oncogene homolog 1 (*Gli1*, hedgehog-responsive gene) and *Axin2* (WNT-responsive gene) in tdTomato+ PDGFRα+ cells compared to tdTomato+ PDGFRα− cells (Fig. 4E). The tdTomato+ PDGFRα− cell population was assumed to be *bona fide* ASMCs (and VSMCs). In line with this assumption, tdTomato+ PDGFRα− cells revealed higher expression levels for *Acta2* and *Lgr6* compared to tdTomato+ PDGFRα+ cells (Fig. 4E). *Lgr6* has been previously shown to identify a subpopulation of WNT-responsive ASMCs that are involved in the repair process following naphthalene injury (Lee et al., 2017). The tdTomato+ PDGFRα− cell population also showed an increase in relative abundance from day 3 to days 7 and 14 (Fig. 4D), indicating that ASMCs also undergo proliferation in response to naphthalene injury, as previously described (Lee et al., 2017; Volckaert et al., 2011).

**Figure 4:**
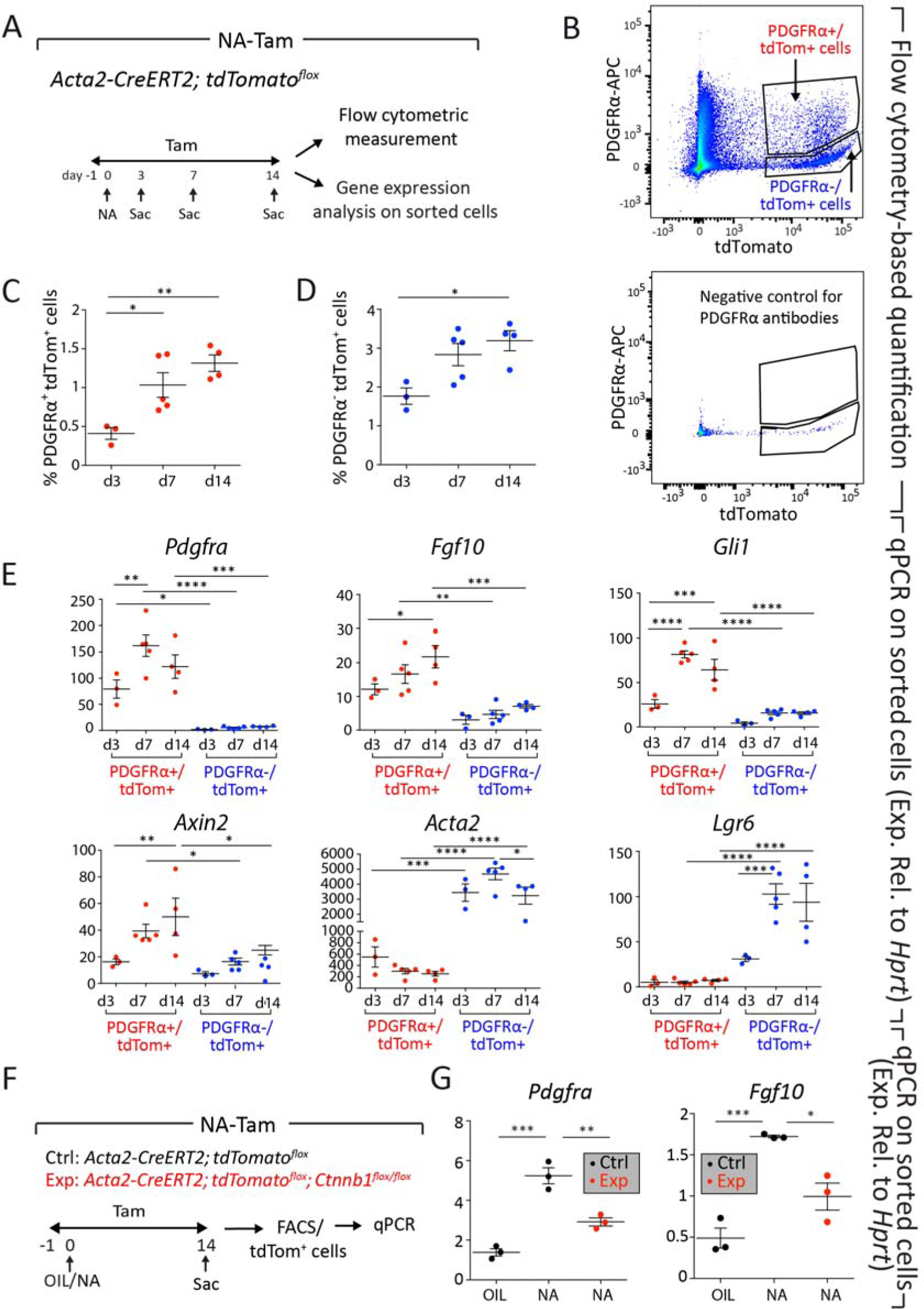
Analysis of the dynamics of the PDGFRα+ tdTomato+ cell population emerging in the NA-Tam group. (A) Experimental setup and time line of naphthalene and tamoxifen treatment. (B) Gating strategy for flow cytometry measurement and sorting of cells based on tdTomato and PDGFRα expression. (C, D) Both PDGFRα+ tdTomato+ cells (red) and PDGFRα− tdTomato+ cells (blue) increase in relative abundance from day 3 to day 14. (E) Quantitative real-time PCR for *Pdgfra, Fgf10, Gli1, Axin2, Acta2* and *Lgr6* using sorted PDGFRα+ tdTomato+ and PDGFRα− tdTomato+ cells. (F) Experimental setup and time line of naphthalene and tamoxifen treatment for analysis of the impact of *Ctnnb1* deletion on inducing *Pdgfra* and *Fgf10* expression. (G) Quantitative real-time PCR for *Pdgfra* and *Fgf10* using sorted tdTomato+ cells. Note that upregulation of *Pdgfra* and *Fgf10* is significantly inhibited in the experimental group. N=3 for d3, N=5 for d7, N=4 for d14 in B-E; N=3 for all experiments in G. * P<0.05, ** P<0.01, *** P<0.001, **** P<0.0001.

As another layer of validation for the RSMC population, *Acta2-CreERT2; tdTomato^flox^*; *Ctnnb1^flox/flox^* (experimental) and *Acta2-CreERT2; tdTomato^flox^* (control) mice were subjected to naphthalene injury and fed tamoxifen-containing pellets up to day 14 (Fig. 4F). Quantitative real-time PCR on sorted cells revealed a 5-fold upregulation of *Pdgfra* and a 3.5-fold upregulation of *Fgf10* in tdTomato+ cells derived from control *Acta2-CreERT2; tdTomato^flox^* mice (Fig. 4G). The induction of these marker genes was significantly attenuated in sorted tdTomato+ cells derived from experimental *Acta2-CreERT2; tdTomato^flox^; Ctnnb1^flox/flox^* mice (Fig. 4G). These results indicate that β-catenin signaling in RSMCs is critical for inducing *Pdgfra* and *Fgf10* expression and consequently epithelial regeneration.

### The GLI1+ lineage contributes to PDGFRα+ RSMCs in the lung

The peribronchial and perivascular regions of the lung are populated by hedgehog-responsive GLI1+ mesenchymal cells (Fig. 5A, B). These cells have been shown to contribute to bleomycin-associated myofibroblasts in lung fibrosis (Kramann et al., 2015) and were also shown to undergo proliferation following naphthalene injury (Peng et al., 2015). Since GLI1+ cells are intermingled with ASMCs in the peribronchial space (Fig. 5A, B), we suspected that the pre-existing GLI1+ pool might serve as a source of RSMCs following naphthalene injury. To test this hypothesis, *Gli1^CreERT2^; tdTomato^flox^* mice were fed tamoxifen-containing pellets for 2 weeks followed by a 2-week washout period and then treated with naphthalene or corn oil (Fig. 5C). This approach allows labeling of pre-existing GLI1+ cells and studying their contribution to epithelial repair and accordingly to RSMC formation. A similar experimental setup was carried out using *Gli1^CreERT2^; tdTomato^flox^; Fgf10^flox/flox^* mice in order to investigate whether FGF10 specifically derived from these cells is required for the repair process (Fig. 5C). Examination of corn oil-treated *Gli1^CreERT2^; tdTomato^flox^* mice revealed significant abundance of tdTomato+ cells around the airways (Fig. 5D, *upper left*). The number of tdTomato+ cells was significantly increased in this region upon naphthalene treatment (Fig. 5D, *upper right*), as previously described (Peng et al., 2015). Strikingly, GLI1+ lineage descendants were also observed in similar regions as *Acta2-CreERT2*-traced RSMCs (Fig. 5F, *upper panel*). Moreover, genetic deletion of both *Fgf10* alleles in pre-existing GLI1+ cells significantly impaired club cell replenishment (Fig. 5D, *lower panel* and Fig. 5E) and yielded less abundance of GLI1+ cells around the airways (Fig. 5F, *lower panel* and Fig. 5G). Quantitative real-time PCR using lung homogenates showed significant downregulation of *Scgb1a1* and *Fgf10* at day 14 after naphthalene treatment in experimental mice compared to controls (Fig. 5H). Furthermore, qPCR on tdTomato+ cells sorted from oil- and naphthalene-treated *Gli1^CreERT2^; tdTomato^flox^* lungs revealed significant downregulation of *Gli1* at day 3 as previously described (Peng et al., 2015), in parallel to a significant, transient upregulation of *Acta2* at day 3 (Fig. 5I). Flow cytometry-based quantification confirmed the increase in the abundance of tdTomato+ ACTA2+ cell population upon naphthalene injury (Fig. 5J). Altogether, these data indicate that the pre-existing GLI1+ cell lineage transiently acquires ACTA2 expression and contributes to RSMC formation during epithelial regeneration.

**Figure 5:**
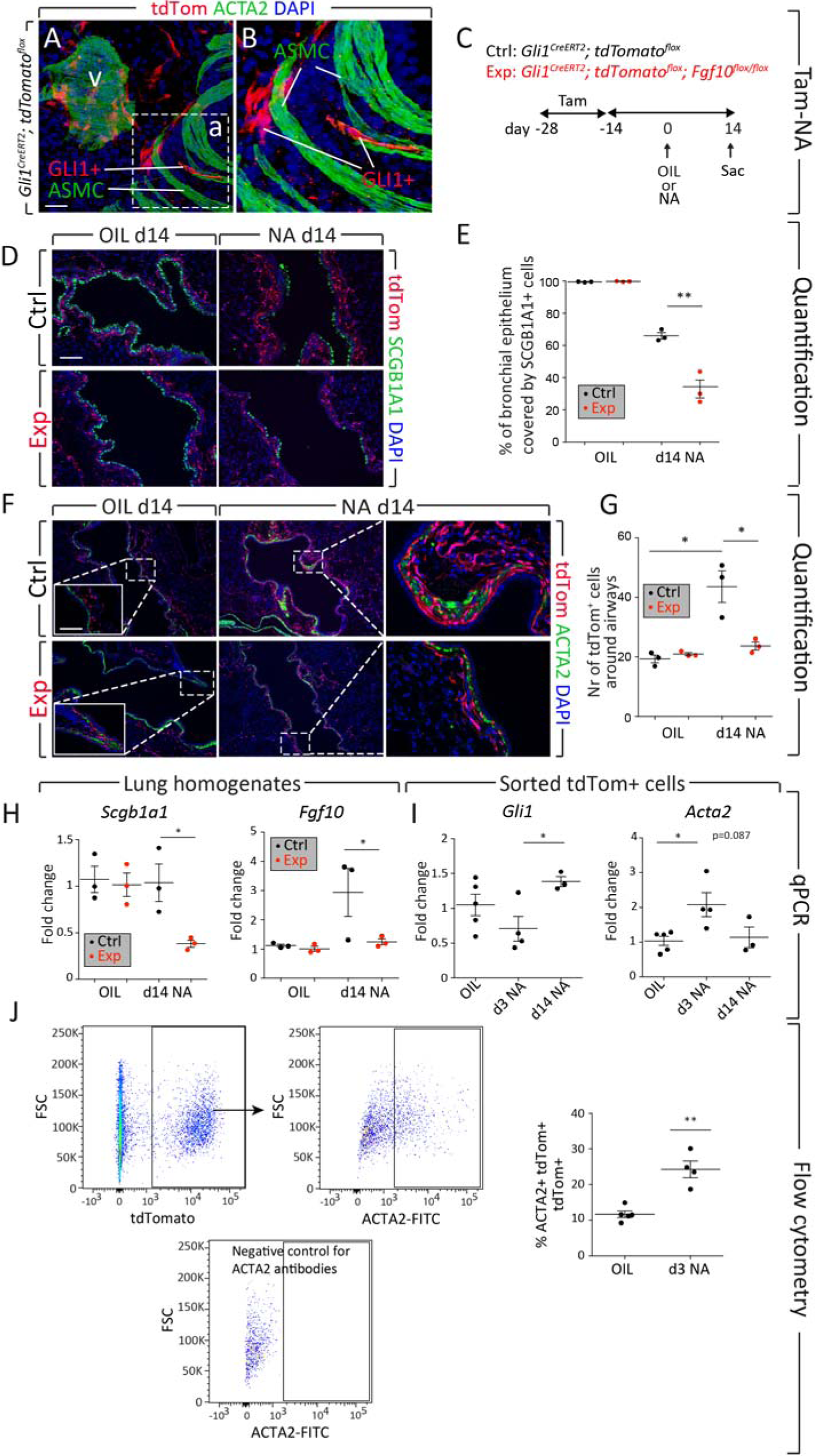
GLI1+ cells give rise to RSMCs and express *Fgf10* that is required for the repair process. (A) Three-dimensional (3D) reconstruction of a *Gli1^CreERT2^; tdTomato^flox^* lung section stained with anti-ACTA2 antibodies. Note the presence of tdTomato+ cells intermingled with the ASMC sheets. The area in the dashed box is magnified in (B). (C) Experimental setup and time line of tamoxifen and naphthalene/corn oil treatment. (D) Immunofluorescence for SCGB1A1 using control and experimental lungs treated with corn oil or naphthalene and analyzed at d14. (E) Quantification of immunofluorescence shown in D demonstrating decreased abundance of SCGB1A1+ cells in the lungs of experimental mice, indicating progressive impairment of epithelial regeneration. (F) Immunofluorescence for ACTA2 and tdTomato using control and experimental lungs. (G) Quantification of the immunofluorescence shown in F. The relative number of tdTomato+ cells around the airways is significantly increased upon naphthalene treatment in control but not in experimental lungs. (H) Quantitative real-time PCR using lung homogenates and showing decreased levels of *Scgb1a1* and *Fgf10* gene expression in experimental mice compared to controls. (I) Quantitative real-time PCR for *Gli1* and *Acta2* using tdTomato+ cells sorted from oil-and naphthalene-treated *Gli1^CreERT2^; tdTomato^flox^* lungs. (J) Gating strategy and quantification of cells based on tdTomato and ACTA2 expression. a – Airway, v – Vessel. Scale bars: 100 μm in D; 100 μm for low magnification and 50 μm for high magnification in F. N=3 for all experiments except for N=2 in d3 NA in (I) and in (J). t-test was used in (J). * P<0.05, ** P<0.01.

### RSMCs support club cell growth in an *in vitro* co-culture organoid system

We then opted to test whether RSMCs possess an intrinsic capacity to support the growth of club cells in a co-culture organoid system. Club cells were sorted using uninjured *Scgb1a1^CreERT2^; tdTomato^flox^* mice and were co-cultured with PDGFRα− tdTomato+ cells or PDGFRα+ tdTomato+ derived from NA-Tam *Acta2-CreERT2; tdTomato^flox^* mice (Fig. 6A). Total tdTomato+ cells from oil-treated *Acta2-CreERT2; tdTomato^flox^* mice were also used as controls. Organoids were allowed to grow for 14 days. This setup allowed comparing club-cell growth-supportive potential of *bona fide* SMCs (PDGFRα− tdTomato+) and RSMCs (PDGFRα+ tdTomato+) derived from naphthalene-injured lungs. The culture conditions used yielded mostly bronchiolospheres (Fig. 6B-D). Strikingly, PDGFRα+ tdTomato+ mesenchymal cells derived from NA-Tam animals yielded higher numbers of organoids (Fig. 6E) with larger diameters (Fig. 6F) compared to PDGFRα− tdTomato+ cells (also derived from NA-Tam animals) and tdTomato+ cells derived from oil-treated controls. Collectively, these data suggest that RSMCs are likely more efficient than ASMCs in supporting the growth of club cells and provide further evidence that these cells are involved in the repair process following naphthalene injury.

**Figure 6:**
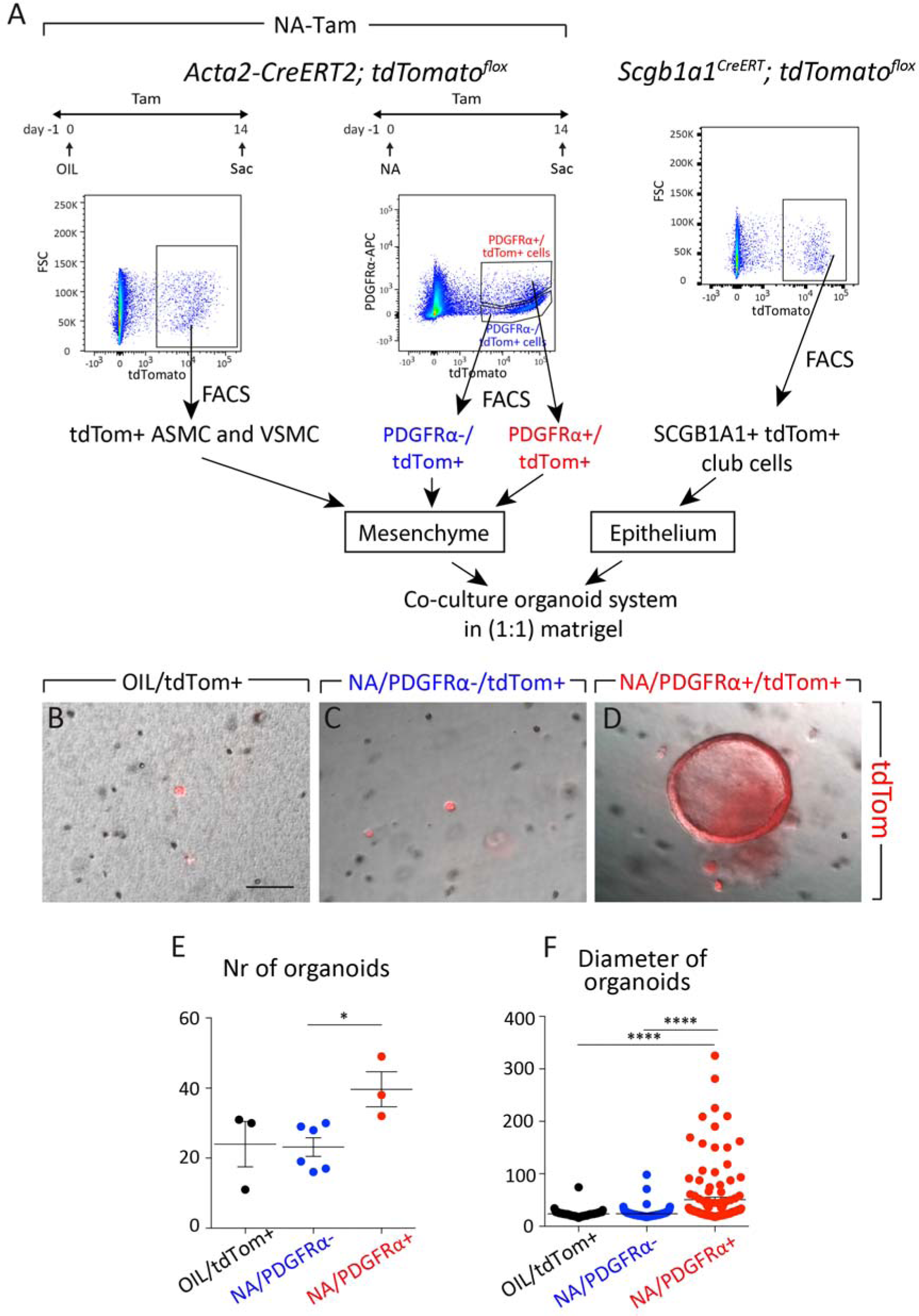
Analysis of the ability of RSMCs to support club cell growth in an *in vitro* co-culture organoid system. (A) Experimental setup for isolating SCGB1A1+ club cells and co-culture with either tdTomato+ cells from oil-treated control lungs, PDGFRα+ tdTomato+ RSMCs or PDGFRα− tdTomato+ cells isolated from naphthalene-treated lungs. (B-D) Imaging of the tdTomato signal in different culture combinations. Note the higher number and bigger diameter of the organoids when SCGB1A1+ cells are cultured with PDGFRα+ tdTomato+ cells (RSMCs) isolated from naphthalene-treated lungs. (E, F) Quantification of the number and diameter of organoids formed under different conditions. Scale bar: 100 μm in B, C, D. N=3 for [OIL/tdTom+] + [SCGB1A1+], N=6 for [NA/PDGFRα− tdTomato+] + [SCGB1A1+], N=3 for [NA/PDGFRα+ tdTomato+] + [SCGB1A1+]. * P<0.05, **** P<0.0001.

## Discussion

Epithelial-mesenchymal interactions represent a hallmark feature of embryonic lung development. These interactions are mediated by diffusible, paracrine-acting growth factors that form gradients in the mesenchyme, thus controlling pattern formation and organogenesis (El Agha and Bellusci, 2014; Morrisey and Hogan, 2010; Shannon and Hyatt, 2004; Warburton et al., 2010). The modes of action of developmental signaling pathways are generally preserved during postnatal life. An exception to this statement is sonic hedgehog signaling that is known to induce mesenchymal proliferation during embryogenesis (Bellusci et al., 1997b) but maintains mesenchymal quiescence during adult life (Peng et al., 2015). Among the best-characterized developmental pathways in the lung are FGF and WNT signaling pathways that are being increasingly implemented in repair processes in the adult lung. In particular, epithelium-derived WNT ligands are emerging as key mediators activating the mesenchymal niche that in turn produces FGF10, leading to epithelial stem-cell activation and regeneration. This mechanism has been shown to be critical for activating at least two types of epithelial stem/progenitor cells in the lung: basal stem cells (Balasooriya et al., 2017; Volckaert et al., 2017) and club cells (Lee et al., 2017; Volckaert et al., 2011).

Despite extensive research, the lineage map of the lung mesenchyme is still poorly understood, with the best-characterized cell types being ASMCs/VSMCs, pericytes, lipofibroblasts (mostly during neonatal stages) and, to a less extent, myofibroblasts. The perivascular region of the lung is populated by GLI1+ mesenchymal cells that constitute a subpopulation of PDGFRβ+ pericytes (Kramann et al., 2015). These cells have been shown to possess mesenchymal stem cell (MSC)-like characteristics such as trilineage differentiation potential *in vitro* (giving rise to adipocytes, osteoblasts and chondrocytes), and also represent a repertoire of myofibroblast progenitors in fibrotic disease (El Agha et al., 2017a; Kramann et al., 2015). On the other hand, the peribronchial domain of the postnatal mouse lung consists of cartilaginous rings (around main-stem bronchi), ASMCs (around conducting airways) and other poorly characterized mesenchymal cells, some of which also express *Gli1*. In this study we demonstrate the complexity of the peribronchiolar region and identify RSMCs as a component of the niche required for activating club cell progenitors. RSMC precursors are intermingled with ASMCs in the confined peribronchiolar space and are activated upon club cell depletion. Our data strongly suggest that these cells originate, at least in part, from GLI1+ peribronchiolar cells. These cells transiently acquire an SMC-like phenotype and contribute to the repair process by producing FGF10 (Fig. 7). Side-by-side comparison of RSMCs (PDGFRα+ tdTomato+) and *bona fide* SMCs (PDGFRα− tdTomato^+^) showed that RSMCs display higher WNT signaling activation (using *Axin2* mRNA levels as readout), higher *Fgf10* expression and greater potential in supporting club cell growth in an *in vitro* co-culture organoid system. However, the caveat here is that the PDGFRα− tdTomato+ population contains a mixture of ASMCs and VSMCs rather than a pure population of ASMCs and this might reduce the colony-forming efficiency and consequently bronchiolosphere number and size.

**Figure 7:**
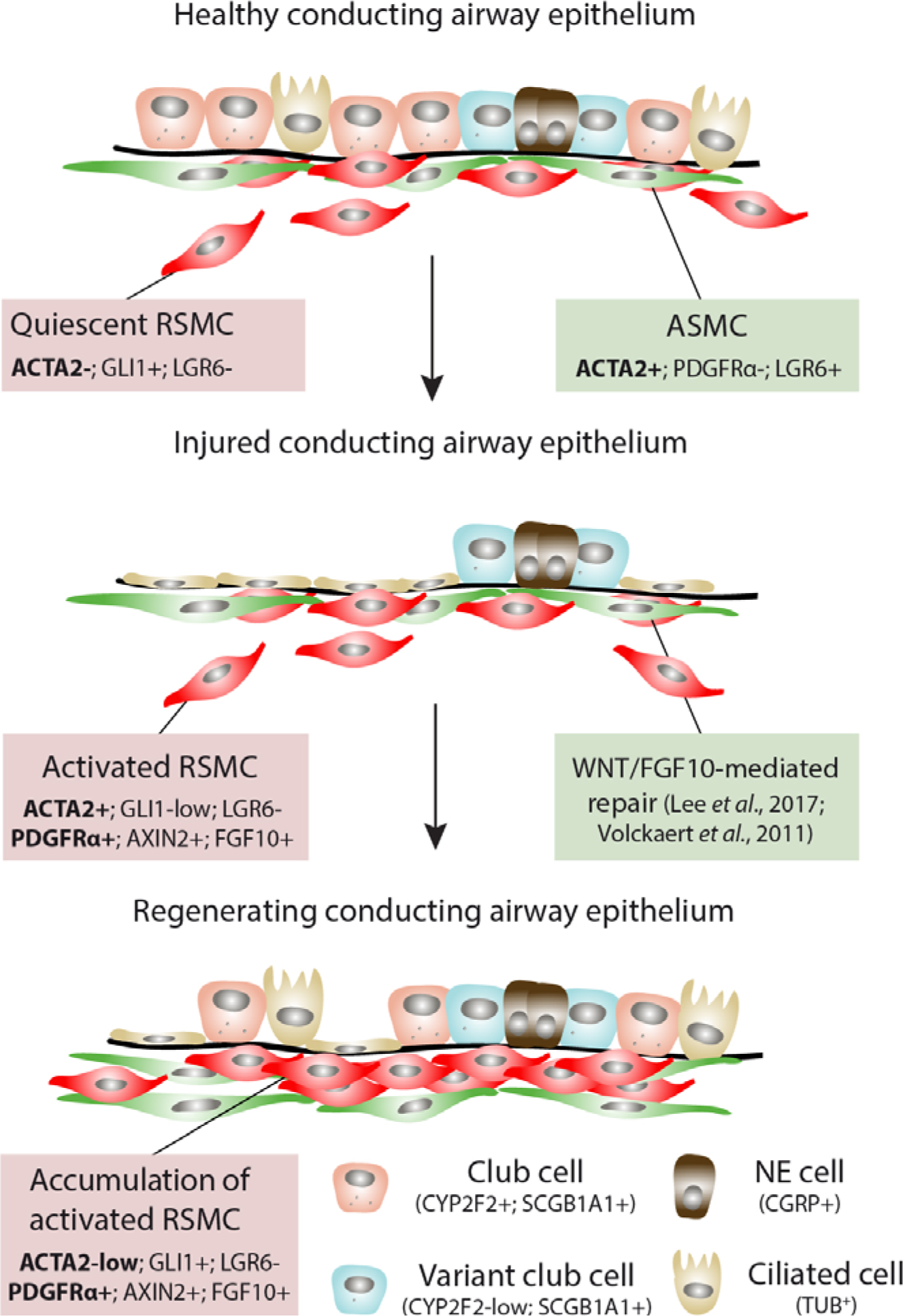
Model summarizing the role of RSMCs as a niche that facilitates epithelial regeneration after naphthalene injury. GLI1+ cells are intermingled with the ASMC layer in the peribronchiolar space. After naphthalene injury, GLI1+ cells transiently acquire ACTA2 expression, give rise to RSMCs (AXIN2+ PDGFRα+) and contribute to epithelial regeneration by producing FGF10. Pre-existing ASMCs have also shown to mediate the repair process via a WNT-FGF10-mediated mechanism (Lee et al., 2017; Volckaert et al., 2011). NE: Neuroendocrine.

A previous study combined single-cell analysis of bulk lung mesenchyme as well as lineage-traced cells and identified a pool of alveolar AXIN2+ PDGFRα+ mesenchymal cells that are involved in type 2 alveolar epithelial cell maintenance as well as a pool of peribronchial AXIN2+ mesenchymal cells (largely PDGFRα−) that give rise to ASMCs upon naphthalene injury (Zepp et al., 2017). Thus, PDGFRα has so far been described as a marker of the (distal) alveolar mesenchymal niche (Barkauskas et al., 2013; Zepp et al., 2017) rather than the (proximal) airway mesenchymal niche. In this study, we identify *Pdgfra* (that is a downstream target of WNT signaling) as a marker of RSMCs that are associated with airway repair. Our data are also in line with the notion that activation of WNT signaling in peribronchial mesenchymal cells promotes epithelial regeneration (Lee et al., 2017; Volckaert et al., 2011; Zepp et al., 2017).

A recent study used scRNA-seq to deconvolute the heterogeneity of lung mesenchyme in response to bleomycin-induced fibrosis (Xie et al., 2018). The authors identified two populations of matrix fibroblasts (COL13A1+ *vs.* COL14A1+) that expanded during fibrosis formation and identified lipofibroblasts as an upstream precursor cell for myofibroblasts, thus confirming our previous report (El Agha et al., 2017b), as well as for matrix fibroblasts (Xie et al., 2018). In the current study, RSMCs display a signature that partially overlaps with COL13A1+ matrix fibroblasts. Whether RSMCs (reported in this study) and COL13A1+ matrix fibroblasts (reported by (Xie et al., 2018)) are entirely distinct populations remains to be determined, especially that different Cre-driver lines, scRNA-seq platforms and other experimental conditions were used in these two independent studies. Accordingly, it remains unclear whether similar mesenchymal subpopulations are activated in response to various models of lung injury.

Our observation that lineage-traced cells were also found in close proximity to bronchiolar epithelial cells raised the possibility that the *Acta2-CreERT2; tdTomato^flox^* mouse line might have labeled surviving progenitor cells undergoing transient epithelial-to-mesenchymal transition prior to regeneration, as previously described (Volckaert et al., 2011). If that scenario was true, tdTomato would have colocalized with the SCGB1A1 stain during the recovery phase at day 14. However, no colocalization between tdTomato and pan-epithelial marker EpCAM or SCGB1A1 was observed at day 14 (Fig. S4), indicating that labeled cells were indeed of mesenchymal origin. The fact that lineage tracing using the *Gli1^CreERT2^* driver, which labels hedgehog-receptive mesenchymal cells, resulted in lineage-labeled cells in a similar anatomical location as RSMCs supports our interpretation. Therefore, our data strongly suggest that the peribronchiolar GLI1+ mesenchymal pool contains RSMC progenitors that are activated and recruited to sites of epithelial injury where they contribute to lung regeneration. A recent report has shown that the GLI1+ cell population overlaps with the PDGFRα+ cell population in the peribronchiolar but not alveolar regions of the adult mouse lung (Wang et al., 2018). Thus, our study complements the growing landscape of knowledge regarding mesenchymal lineage formation and behavior during repair after injury, and warrants further research on cellular and molecular mechanisms involved in many life-threatening and chronic respiratory diseases such as asthma and chronic obstructive pulmonary disease.

## Acknowledgements

E.E.A. was funded by grants from the Excellence Cluster Cardio-Pulmonary System (ECCPS) and University Hospital Giessen and Marburg (UKGM). E.E.A. also acknowledges the support of the German Center for Lung Research (DZL) and the Cardio-Pulmonary Institute (CPI, EXC 2026, Project ID: 390649896). S.B. was supported by grants from the Deutsche Forschungsgemeinschaft (DFG; BE4443/14-1, BE4443/6-1, KFO309 P7 and SFB CRC1213-projects A02 and A04), the DZL and the First Affiliated Hospital of Wenzhou Medical university. S.B. also acknowledges the support of the CPI. J.S.Z was funded through a start-up package from Wenzhou Medical University and the National Natural Science Foundation of China (grant number 81472601). S.H. was supported by the UKGM (FOKOOPV), the DZL and grants from the DFG (KFO309 P2/8; SFB1021 C05, SFB TR84 B9). We would also like to thank Dr. Athanasios Fysikopoulos and Prof. Christos Samakovlis for providing the *Scgb1a1^CreERT2^; tdTomato^flox^* mice used for the organoid experiments. We also thank Irina Shalashova for her help in obtaining tile scans of lung sections.

---

**Supplementary Figure S1:**
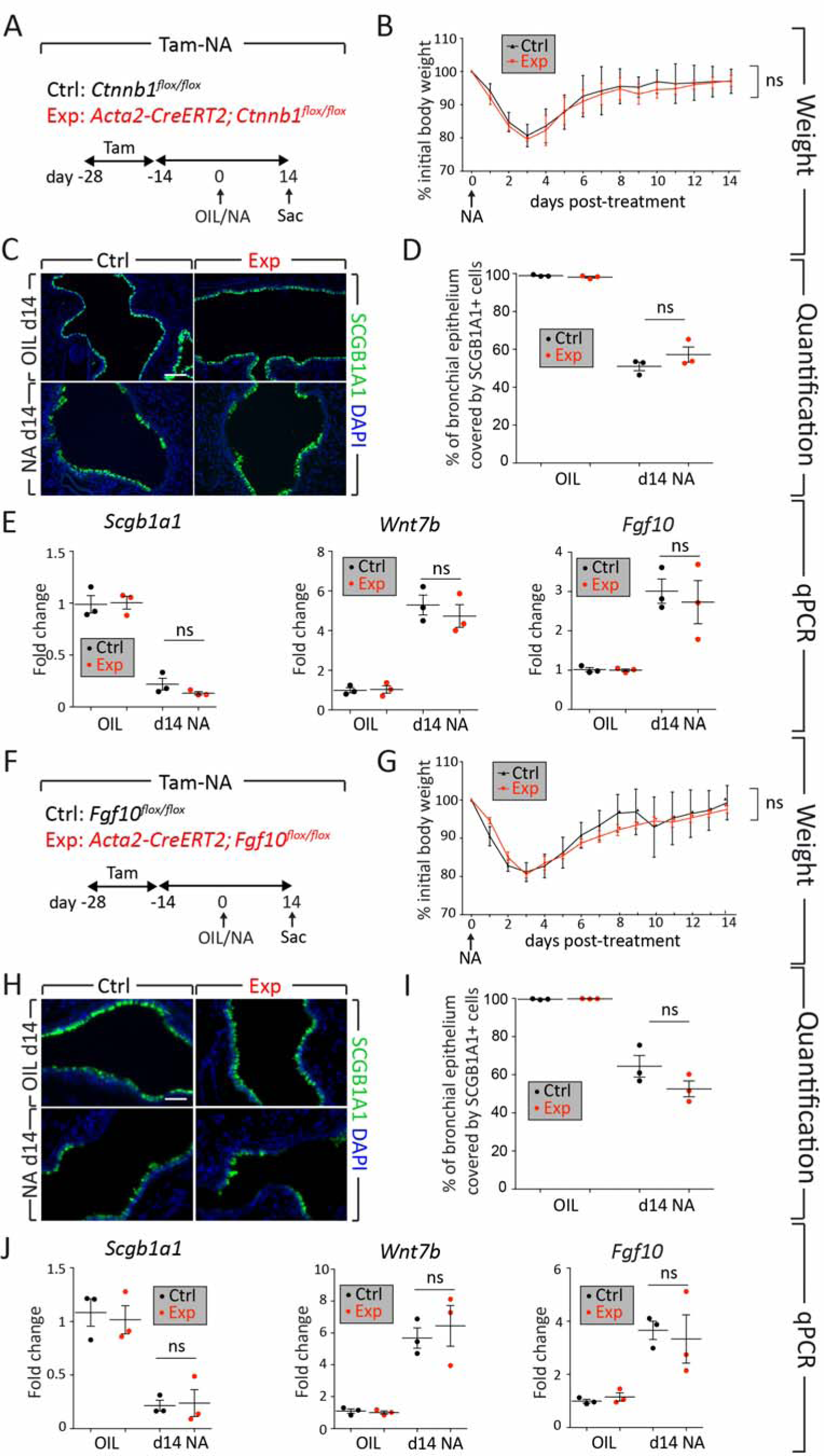
Deletion of *Ctnnb1* or *Fgf10* in ACTA2+ cells before naphthalene injury does not affect epithelial regeneration. (A, F) Experimental setup and time line of tamoxifen and naphthalene treatment. (B, G) Weight loss curve showing no difference between experimental and control mice. (C, H) Immunofluorescence for SCGB1A1 using control and experimental lungs. (D, I) Quantification of immunofluorescence demonstrating no difference in the percentage of airway epithelium covered by SCGB1A1+ cells. (E, J) Quantitative real-time PCR using lung homogenates and showing no difference in the expression levels of *Scgb1a1*, *Wnt7b* and *Fgf10* between control and experimental mice. ns – Not significant. Scale bars: 50 μm in C; 100 μm in H. N=3 for all experiments.

**Supplementary Figure S2:**
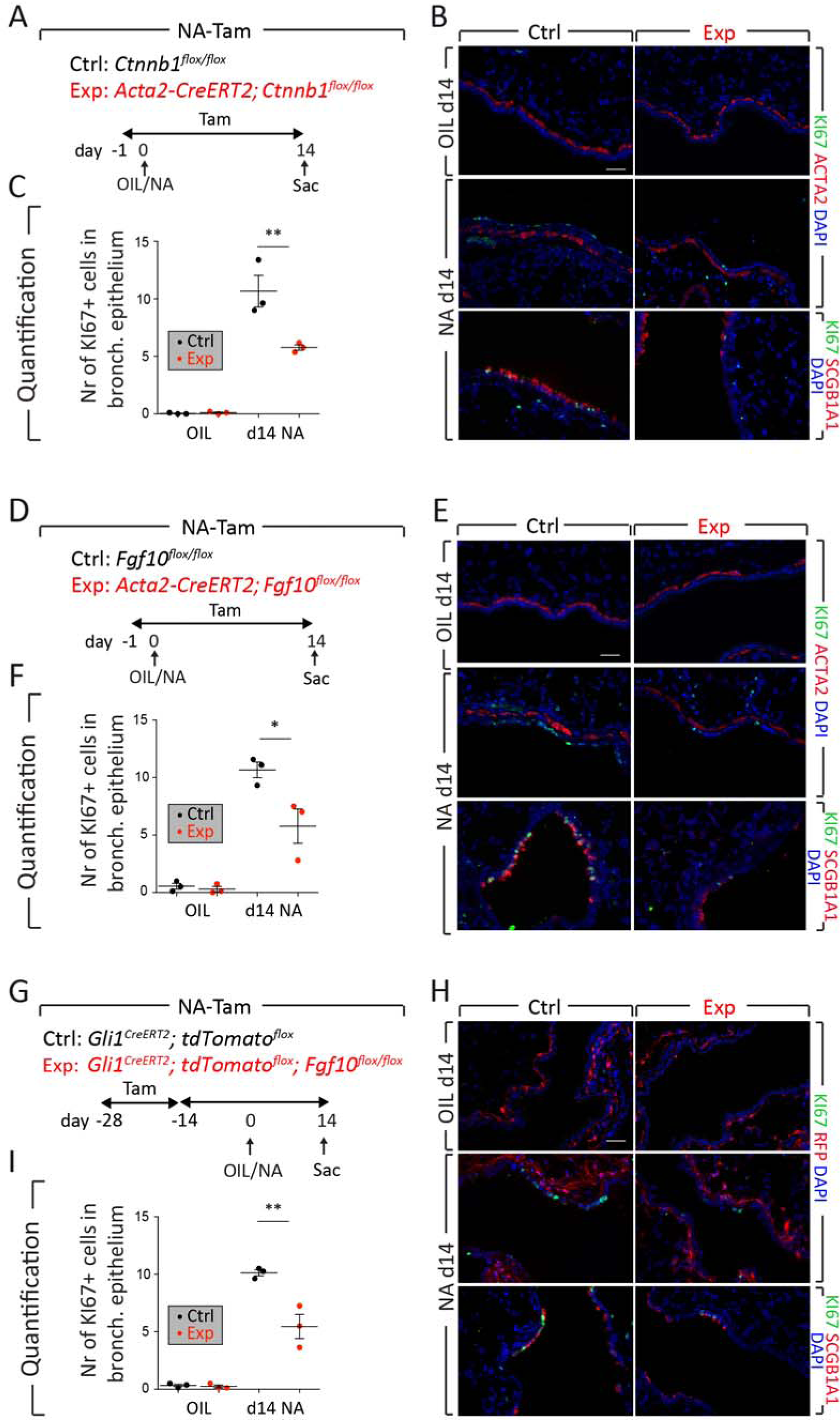
Analysis of proliferation after naphthalene injury. (A, D, G) Experimental setup and time line of tamoxifen and naphthalene/corn oil treatment. (B, E, H) Immunofluorescence for KI67, SCGB1A1, ACTA2 and RFP at d14. (C, F, I) Quantification of KI67+ cells in the bronchiolar epithelium. Scale bar: 35 μm. N=3. * P<0.05, ** P<0.01.

**Supplementary Figure S3:**
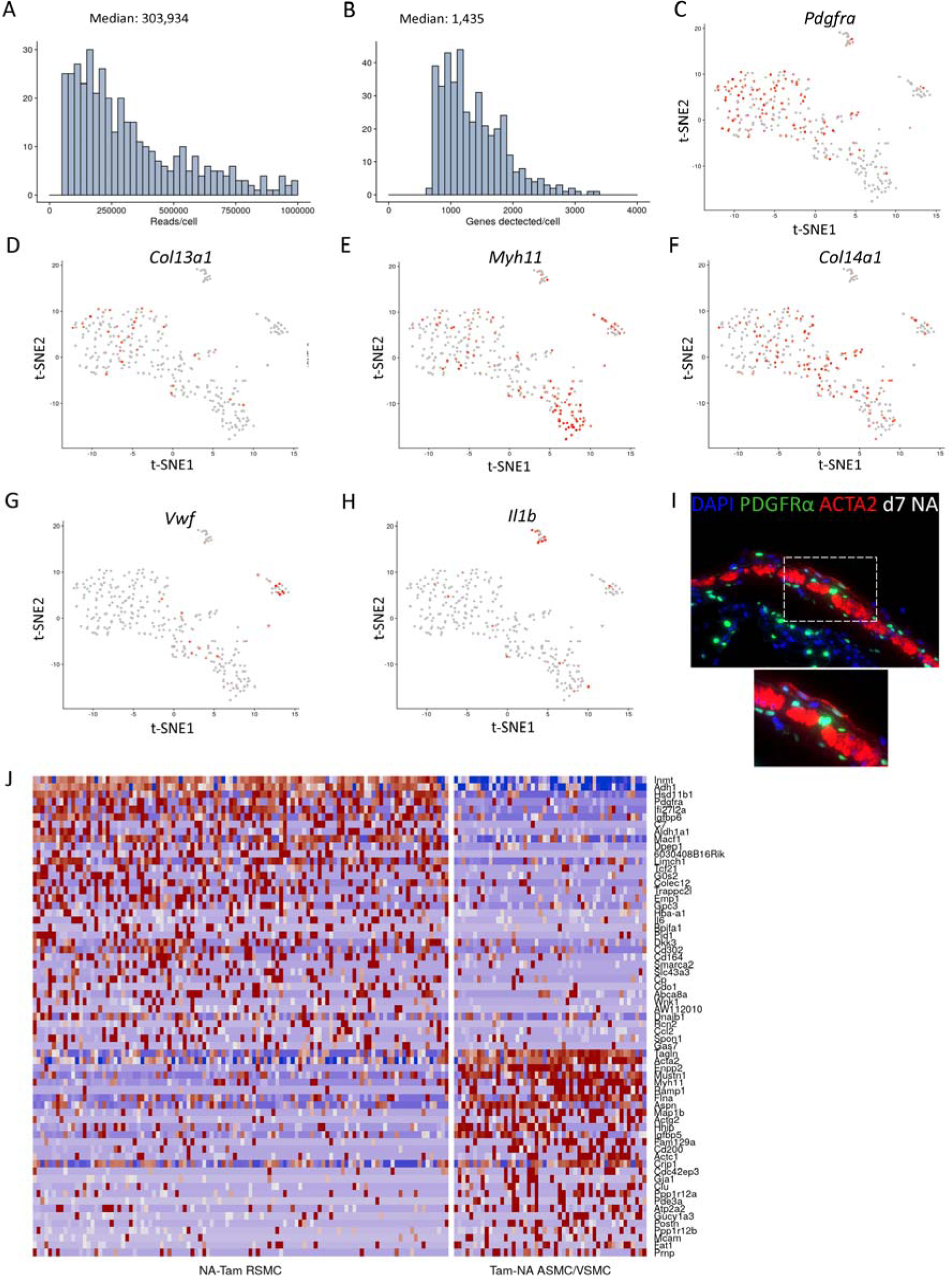
Quality control for the scRNA-seq experiment. (A, B) Median number of reads per cell and number of genes detected per cell are shown. (C-G) t-SNE plots showing enrichment of selected markers in various cellular clusters. (H) Heat map depicting top regulated genes between NA-Tam RSMCs and Tam-NA ASMC/VSMC. (I) Immunofluorescence for ACTA2 on a *Pdgfra^H2B-EGFP^* mouse lung at day 7 after naphthalene injury. The area in the box is magnified in the lower panel. Note the presence of ACTA2-low EGFP+ cells in the peribronchiolar space.

**Supplementary Figure S4:**
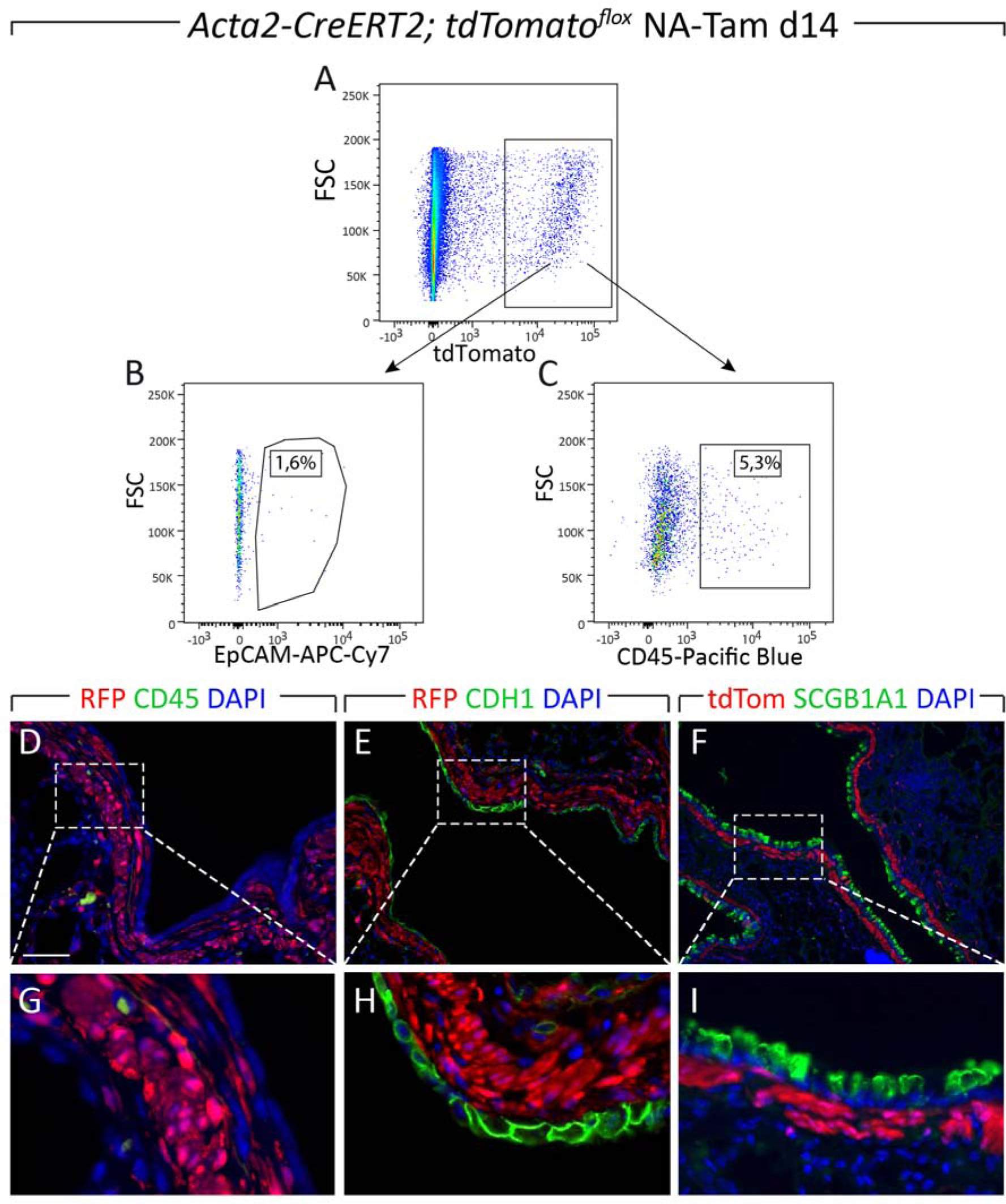
ACTA2+ cells labeled after injury do not express epithelial or leukocyte markers at d14. (A-C) Flow cytometry plots showing negligible overlap between tdTomato+ cells and EpCAM+ epithelial cells (1,6%) or CD45+ cells leukocytes (5,3%) in lungs of NA-Tam *Acta2-CreERT2; tdTomato^flox^* mice. **(D-I)** Immunofluorescence for CD45, CDH1 (E-CAD), SCGB1A1 and tdTomato. (G-I) High magnification of the boxes shown in D-F. Scale bar: 50 μm. N=3 for all experiments.

## References

Abler, L.L., Mansour, S.L., and Sun, X. (2009). Conditional gene inactivation reveals roles for Fgf10 and Fgfr2 in establishing a normal pattern of epithelial branching in the mouse lung. Dev Dyn 238, 1999–2013.

El Agha, E., and Bellusci, S. (2014). Walking along the Fibroblast Growth Factor 10 Route: A Key Pathway to Understand the Control and Regulation of Epithelial and Mesenchymal Cell-Lineage Formation during Lung Development and Repair after Injury. Scientifica (Cairo). 2014, 538379.

El Agha, E., Herold, S., Al Alam, D., Quantius, J., MacKenzie, B., Carraro, G., Moiseenko, A., Chao, C.-M., Minoo, P., Seeger, W., et al. (2014). Fgf10-positive cells represent a progenitor cell population during lung development and postnatally. Development 141, 296–306.

El Agha, E., Kramann, R., Schneider, R.K., Li, X., Seeger, W., Humphreys, B.D., and Bellusci, S. (2017a). Mesenchymal Stem Cells in Fibrotic Disease. Cell Stem Cell 21, 166–177.

El Agha, E., Moiseenko, A., Kheirollahi, V., De Langhe, S., Crnkovic, S., Kwapiszewska, G., Szibor, M., Kosanovic, D., Schwind, F., Schermuly, R.T., et al. (2017b). Two-Way Conversion between Lipogenic and Myogenic Fibroblastic Phenotypes Marks the Progression and Resolution of Lung Fibrosis. Cell Stem Cell 20, 261–273.e3.

Balasooriya, G.I., Goschorska, M., Piddini, E., and Rawlins, E.L. (2017). FGFR2 is required for airway basal cell self-renewal and terminal differentiation. Development 144, 1600–1606.

Barkauskas, C.E., Cronce, M.J., Rackley, C.R., Bowie, E.J., Keene, D.R., Stripp, B.R., Randell, S.H., Noble, P.W., and Hogan, B.L.M. (2013). Type 2 alveolar cells are stem cells in adult lung. J. Clin. Invest. 123, 3025–3036.

Bellusci, S., Grindley, J., Emoto, H., Itoh, N., and Hogan, B.L. (1997a). Fibroblast growth factor 10 (FGF10) and branching morphogenesis in the embryonic mouse lung. Development 124, 4867–4878.

Bellusci, S., Furuta, Y., Rush, M.G., Henderson, R., Winnier, G., and Hogan, B.L. (1997b). Involvement of Sonic hedgehog (Shh) in mouse embryonic lung growth and morphogenesis. Development 124, 53–63.

Boyd, M.R. (1977). Evidence for the Clara cell as a site of cytochrome P450-dependent mixed-function oxidase activity in lung. Nature 269, 713–715.

Butler, A., Hoffman, P., Smibert, P., Papalexi, E., and Satija, R. (2018). Integrating single-cell transcriptomic data across different conditions, technologies, and species. Nat. Biotechnol. 36, 411–420.

Clara, M. (1937). Zur histobiologie des bronchalepithels. Z. Mikrosk. Anat. Forsch. 41, 321–347.

Cohen, E.D., Ihida-Stansbury, K., Lu, M.M., Panettieri, R.A., Jones, P.L., and Morrisey, E.E. (2009). Wnt signaling regulates smooth muscle precursor development in the mouse lung via a tenascin C/PDGFR pathway. J. Clin. Invest. 119, 2538–2549.

Fanucchi, M. V, Murphy, M.E., Buckpitt, A.R., Philpot, R.M., and Plopper, C.G. (1997). Pulmonary cytochrome P450 monooxygenase and Clara cell differentiation in mice. Am J Respir Cell Mol Biol 17, 302–314.

Giangreco, A., Reynolds, S.D., and Stripp, B.R. (2002). Terminal bronchioles harbor a unique airway stem cell population that localizes to the bronchoalveolar duct junctionGiangreco, A., Reynolds, S.D., and Stripp, B.R. (2002). Terminal bronchioles harbor a unique airway stem cell population that localizes to Am J Pathol 161, 173–182.

Hong, K.U., Reynolds, S.D., Giangreco, A., Hurley, C.M., and Stripp, B.R. (2001). Clara Cell Secretory Protein–Expressing Cells of the Airway Neuroepithelial Body Microenvironment Include a Label-Retaining Subset and Are Critical for Epithelial Renewal after Progenitor Cell Depletion. Am. J. Respir. Cell Mol. Biol. 24, 671–681.

Kramann, R., Schneider, R.K., DiRocco, D.P., Machado, F., Fleig, S., Bondzie, P.A., Henderson, J.M., Ebert, B.L., and Humphreys, B.D. (2015). Perivascular Gli1+ Progenitors Are Key Contributors to Injury-Induced Organ Fibrosis. Cell Stem Cell 16, 51–66.

Lee, J.H., Tammela, T., Hofree, M., Choi, J., Marjanovic, N.D., Han, S., Canner, D., Wu, K., Paschini, M., Bhang, D.H., et al. (2017). Anatomically and Functionally Distinct Lung Mesenchymal Populations Marked by Lgr5 and Lgr6. Cell 170, 1149–1163.e12.

Macosko, E.Z., Basu, A., Satija, R., Nemesh, J., Shekhar, K., Goldman, M., Tirosh, I., Bialas, A.R., Kamitaki, N., Martersteck, E.M., et al. (2015). Highly Parallel Genome-wide Expression Profiling of Individual Cells Using Nanoliter Droplets. Cell 161, 1202–1214.

Mahvi, D., Bank, H., and Harley, R. (1977). Morphology of a naphthalene-induced bronchiolar lesion. Am. J. Pathol. 86, 558–572.

Mailleux, A.A., Kelly, R., Veltmaat, J.M., De Langhe, S.P., Zaffran, S., Thiery, J.P., and Bellusci, S. (2005). Fgf10 expression identifies parabronchial smooth muscle cell progenitors and is required for their entry into the smooth muscle cell lineage. Development 132, 2157–2166.

Morrisey, E.E., and Hogan, B.L. (2010). Preparing for the first breath: genetic and cellular mechanisms in lung development. Dev Cell 18, 8–23.

Peng, T., Frank, D.B., Kadzik, R.S., Morley, M.P., Rathi, K.S., Wang, T., Zhou, S., Cheng, L., Lu, M.M., and Morrisey, E.E. (2015). Hedgehog actively maintains adult lung quiescence and regulates repair and regeneration. Nature 526, 578–582.

Ramasamy, S.K., Mailleux, A.A., Gupte, V. V, Mata, F., Sala, F.G., Veltmaat, J.M., Del Moral, P.M., De Langhe, S., Parsa, S., Kelly, L.K., et al. (2007). Fgf10 dosage is critical for the amplification of epithelial cell progenitors and for the formation of multiple mesenchymal lineages during lung development. Dev Biol 307, 237–247.

Rawlins, E.L., Okubo, T., Xue, Y., Brass, D.M., Auten, R.L., Hasegawa, H., Wang, F., and Hogan, B.L.M. (2009). The role of Scgb1a1+ Clara cells in the long-term maintenance and repair of lung airway, but not alveolar, epithelium. Cell Stem Cell 4, 525–534.

Reynolds, S.D., Giangreco, A., Power, J.H.T., and Stripp, B.R. (2000). Neuroepithelial Bodies of Pulmonary Airways Serve as a Reservoir of Progenitor Cells Capable of Epithelial Regeneration. Am. J. Pathol. 156, 269–278.

Sekine, K., Ohuchi, H., Fujiwara, M., Yamasaki, M., Yoshizawa, T., Sato, T., Yagishita, N., Matsui, D., Koga, Y., Itoh, N., et al. (1999). Fgf10 is essential for limb and lung formation. Nat Genet 21, 138–141.

Shannon, J.M., and Hyatt, B. a (2004). Epithelial-mesenchymal interactions in the developing lung. Annu. Rev. Physiol. 66, 625–645.

Stripp, B.R., Maxson, K., Mera, R., and Singh, G. (1995). Plasticity of airway cell proliferation and gene expression after acute naphthalene injury. Am J Physiol 269, L791–9.

Urness, L.D., Paxton, C.N., Wang, X., Schoenwolf, G.C., and Mansour, S.L. (2010). FGF signaling regulates otic placode induction and refinement by controlling both ectodermal target genes and hindbrain Wnt8a. Dev Biol 340, 595–604.

Volckaert, T., Dill, E., Campbell, A., Tiozzo, C., Majka, S., Bellusci, S., and De Langhe, S.P. (2011). Parabronchial smooth muscle constitutes an airway epithelial stem cell niche in the mouse lung after injury. J Clin Invest 121, 4409–4419.

Volckaert, T., Yuan, T., Chao, C.-M., Bell, H., Sitaula, A., Szimmtenings, L., El Agha, E., Chanda, D., Majka, S., Bellusci, S., et al. (2017). Fgf10-Hippo Epithelial-Mesenchymal Crosstalk Maintains and Recruits Lung Basal Stem Cells. Dev. Cell 43, 48–59.e5.

Wang, C., Mochel, N.S.R. de, Christenson, S.A., Cassandras, M., Moon, R., Brumwell, A.N., Byrnes, L.E., Li, A., Yokosaki, Y., Shan, P., et al. (2018). Expansion of hedgehog disrupts mesenchymal identity and induces emphysema phenotype. J. Clin. Invest. 128.

Warburton, D., El-Hashash, A., Carraro, G., Tiozzo, C., Sala, F., Rogers, O., De Langhe, S., Kemp, P.J., Riccardi, D., Torday, J., et al. (2010). Lung organogenesis. Curr. Top. Dev. Biol. 90, 73–158.

Wendling, O., Bornert, J.-M., Chambon, P., and Metzger, D. (2009). Efficient temporally-controlled targeted mutagenesis in smooth muscle cells of the adult mouse. Genesis 47, 14–18.

Winkelmann, A., and Noack, T. (2010). The Clara cell: A “Third Reich eponym”? Eur. Respir. J. 36, 722–727.

Xie, T., Wang, Y., Deng, N., Liang, J., Noble, P.W., Xie, T., Wang, Y., Deng, N., Huang, G., Taghavifar, F., et al. (2018). Single-Cell Deconvolution of Fibroblast Heterogeneity in Mouse Pulmonary Fibrosis Article Single-Cell Deconvolution of Fibroblast Heterogeneity in Mouse Pulmonary Fibrosis. Cell Rep. 22, 3625–3640.

Zepp, J.A., Zacharias, W.J., Frank, D.B., Cavanaugh, C.A., Zhou, S., Morley, M.P., and Morrisey, E.E. (2017). Distinct Mesenchymal Lineages and Niches Promote Epithelial Self-Renewal and Myofibrogenesis in the Lung. Cell 170, 1134–1148.e10.

